# Mesoscopic whole-brain T_2_*-weighted and associated quantitative MRI in healthy humans at 10.5 T

**DOI:** 10.1101/2025.04.21.649819

**Authors:** Jiaen Liu, Peter van Gelderen, Jacco A. de Zwart, Jeff H. Duyn, Yujia Huang, Andrea Grant, Edward Auerbach, Matt Waks, Russell Lagore, Lance Delabarre, Alireza Sadeghi-Tarakameh, Yigitcan Eryaman, Gregor Adriany, Kamil Ugurbil, Xiaoping Wu

**Affiliations:** Advanced Imaging Research Center, UT Southwestern Medical Center, Dallas, TX, USA; Radiology, UT Southwestern Medical Center, Dallas, TX, USA; Advanced MRI section, NINDS, NIH, Bethesda, MD, USA; Center for Magnetic Resonance Research, Radiology, Medical School, University of Minnesota Twin Cities, Minneapolis, MN, USA

**Keywords:** ultrahigh field MRI, 10.5 T, T_2_*-weighted MRI, high-density RF coil, mesoscopic MRI

## Abstract

**Purpose:** To demonstrate the feasibility and performance of mesoscopic whole brain T_2_*-weighted (T_2_*w) MRI at 10.5 T by combining a motion-robust multi-echo gradient-echo (GRE) method with high-density RF receive arrays.

**Methods:** Multi-echo GRE data were collected in healthy adults at isotropic 0.5 mm resolution using a custom-built 16-channel transmit/80-channel receive (16Tx/80Rx) RF coil. Whole brain images were reconstructed with navigator-guided joint motion and field correction and were used for quantitative *R_2_*\* and susceptibility (*χ*) mapping. Intrinsic signal-to-noise ratio (iSNR) and quantification precision for *R_2_*\* and *χ* were also estimated. The results were compared with those obtained in the same subjects with matched resolution at 7 T using the commercial Nova 1Tx/32Rx coil, to demonstrate the iSNR and quantification precision gains at 10.5 T. G-factors were also calculated at each field strength to evaluate parallel imaging performances. To demonstrate the benefit of increased parallel imaging performances at 10.5 T, whole brain images with higher acceleration were also obtained using a custom-built 16Tx/128Rx coil.

**Results:** the utilized motion robust GRE sequence and reconstruction effectively reduced artifacts from motion and field changes during scans, producing high-quality whole-brain T_2_*w images and multi-parametric maps at 10.5 T with delineation of fine-scale brain structures. Compared to 7 T, the 10.5 T approach led to gains in both iSNR and quantification precision of *R_2_*\* and *χ*. Quantitatively, iSNR estimated in the peripheral cortical gray matter increased by 42%. Parallel imaging performances were also improved at 10.5 T owing to the utilized high-density coils compared to the commonly used commercially available coil at 7 T, allowing high-quality images with up to 12-fold combined acceleration when using the 128Rx coil.

**Conclusion:** It is feasible to perform motion-robust whole-brain mesoscopic multi-echo gradient echo imaging of the human brain at 10.5 T. Intrinsic SNR and quantification precision of *R*_2_* and *χ* were estimated and compared with 7 T results. The results presented here may shed light on future optimal implementation of anatomic T_2_*w brain MRI at ultrahigh field beyond 7 T.

## 1. Introduction

The pursuit of mesoscopic resolution in human brain imaging (Edwards et al., 2018), bridging the gap between macroscopic anatomical visualization and microscopic cellular details, is a frontier in magnetic resonance imaging (MRI). Ultra-high-field (UHF) MRI systems operating at 7 tesla (T) and above unlock unprecedented signal-to-noise and contrast-to-noise ratios (SNR and CNR) (Duyn, 2012; Duyn et al., 2007; Le Ster et al., 2022; Pohmann et al., 2016; Ugurbil, 2014; Uğurbil, 2018; Waks et al., 2025; Lagore et al., 2025), enabling finer spatial resolution and enhanced sensitivity to mesoscopic tissue structures. Leveraging differences in tissue magnetic susceptibility (*χ*) to reveal intricate brain structures such as cortical layers (Duyn et al., 2007; Fukunaga et al., 2010; van Gelderen et al., 2023), microvasculature (Shen et al., 2020) and iron-rich subcortical nuclei (Bianciardi et al., 2015; Duyn et al., 2007), susceptibility-weighted imaging, also called “T_2_*-weighted (T_2_*w) imaging”, particularly benefits from UHF MRI due to the synergistically enhanced signal-to-noise ratio (SNR) and susceptibility-induced contrast with increasing field strength. When acquired with multi-echo gradient echo (ME-GRE) sequences, this modality along with its derived quantitative measurements, such as the transverse relaxation rate (*R_2_\**, equal to *1/T_2_\**) and magnetic susceptibility *χ*, offers a window for studying brain anatomy (Duyn et al., 2007) and pathology (Bartolini et al., 2019; Liu et al., 2021; van Rooden et al., 2014) in vivo at a spatial resolution previously attainable only ex vivo or with impractically long scan times (Lüsebrink et al., 2017).

Human brain MRI at 10.5 T was demonstrated to be feasible (Sadeghi-Tarakameh et al., 2020), safe (Grant et al., 2020) and capable of delivering high SNR and parallel imaging performance with high-density radiofrequency (RF) receive arrays (Lagore et al., 2025; Waks et al., 2025). It holds promises to further improve T_2_*w imaging resolution beyond what has been achieved at 7 T (Zwanenburg et al., 2011; Gulban et al., 2022; van Gelderen et al., 2023). However, at 10.5 T, challenges related to subject motion, motion-related magnetic field (B_0_) fluctuations (Liu et al., 2018), B_0_, and RF (B_1_) field inhomogeneities are further amplified. In addition, there is a lack of literature documenting T_2_*w contrast increase at 10.5 T and beyond, obscuring the optimal design of T_2_*w sequence parameters for realizing the full benefits of 10.5 T T_2_*w MRI.

In this work, we demonstrate the feasibility of mesoscopic whole-brain T_2_*w anatomical MRI in healthy humans at 10.5 T and systematically evaluate the improvement of *R_2_\** and magnetic susceptibility *χ* quantification precision compared to 7 T. Central to the approach was the use of a motion-robust 3D ME-GRE sequence paired with custom-built high-density RF receive arrays. The motion robustness was achieved through a retrospective navigator-guided joint correction framework (Liu et al., 2020) to simultaneously address rigid-body head motion and motion-induced B_0_ field changes, as developed in our previous work (Liu et al., 2020; van Gelderen et al., 2023). Two high-density head RF receive arrays approved by the U.S. Food and Drug Administration (Lagore et al., 2025; Waks et al., 2025), optimized for SNR and parallel imaging efficiency, were utilized to facilitate high resolution imaging across the entire brain within a clinically relevant scan time. This work presents the initial results of T_2_*w MRI and associated quantitative mapping of *R_2_\** and *χ* at 10.5 T, showcasing detailed anatomic contrast, efficacy of the joint motion and field correction strategy and increased precision and parallel imaging capability. The results establish a practical foundation for pushing the boundaries of in vivo brain imaging based on T_2_*w MRI at 10.5 T, with the potential to advance our understanding of neuroanatomy with unprecedented details.

## 2. Methods

### 2.1 MRI experiments

Two healthy human volunteers (1 male and 1 female) were recruited and scanned in the MRI experiments after having provided a signed informed consent form approved by the local Institutional Review Board. Data were first acquired on a Siemens Magnetom 10.5 T MR scanner (Siemens Healthineers, Erlangen, Germany) equipped with 16-channel RF transmission and 128-channel signal reception systems. Two custom-built high-density RF coils were utilized for data collection: one with 80 receive channels (80Rx) (Waks et al., 2025) and another 128 receive channels (128Rx) (Lagore et al., 2025). The 128Rx coil was included to demonstrate the parallel imaging performance at 10.5 T. Both coils included 16 independent transmit channels for parallel transmission. However, in this work, the transmit channels operated in the circularly polarized (CP) mode for RF transmission to mimic a single-channel transmit setup. For comparison, the same human volunteers were scanned at 7 T on a Siemens Terra MR scanner (Siemens Healthineers, Erlangen, Germany). The commercial Nova single-channel transmit 32-channel receive (1Tx/32Rx) RF coil (Nova Medical, Wilmington, MA, USA) was utilized for data collection, so as to compare to industry standard scanning available at all 7 T sites. Both the 10.5 T and 7 T MR systems were equipped with the SC72 whole-body gradient system with a 70 mT/m maximal strength and a 200 T/m/s slew rate.

Experiments were first performed using the 16Tx/80Rx coil to demonstrate the utility of the adopted acquisition and correction strategies for whole brain imaging at 10.5 T. For each volunteer, a 3D T_2_*w ME-GRE sequence with embedded volumetric navigators was used for data acquisition. Relevant imaging parameters included isotropic 0.5 mm resolution, 240×180×128 mm^3^ field of view (FOV), 35 ms repetition time (TR), 4 echo times (TEs), TE_1_ of 10.2 ms, echo spacing (ES) of 4.9 ms, 12° nominal flip angle, 208 Hz/pixel readout receiver bandwidth and 2×3 2D parallel imaging acceleration with controlled aliasing for parallel imaging (CAIPI) (Breuer et al., 2006). The 2D acceleration was applied in the phase-encoding (left-right) and slice-encoding (head-foot) directions, leading to a total scan time of ∼13.3 min.

The embedded volumetric navigator images meant for real-time motion and B_0_ measurement were collected during the time before the first echo using a multi-shot 3D echo-planar imaging (EPI) method as described in our previous work (Liu et al., 2020; van Gelderen et al., 2023). Specifically, the navigator images were acquired with 5×5.6×8 mm^3^ spatial resolution, 48×32×24 matrix size, 4.3 ms TE and 4×2 acceleration with blipped CAIPI (Poser et al., 2014). An accelerated navigator image volume was acquired in 12 TRs (420 ms).

To evaluate SNR and quantification precision improvement of 10.5 T vs. 7 T, whole-brain data at 7 T was also collected in the same two volunteers with matched imaging parameters including FOV, resolution, 2D acceleration, TE_1_ and ES. However, due to the longer T_2_* values, the 7 T data collection opted for a 6-echo protocol, leading to a longer TR of 45 ms, a higher nominal flip angle of 14°, and a longer total scan time of ∼17.2 min. The embedded volumetric navigator images were collected using the same imaging parameters as in 10.5 T data acquisition, except that the temporal acquisition interval was increased to 540 ms due to the increased TR.

At both field strengths, fully sampled calibration data aimed at coil sensitivity mapping needed for image reconstruction were acquired in a separate reference scan using a multi-slice 2D GRE sequence. Specifically, multi-slice 2D GRE images with whole brain coverage were obtained with the following relevant imaging parameters: 4 mm isotropic in-plane resolution, 256×192 mm^2^ FOV, 4 mm slice thickness and 50 axial slices. At each field strength, TE was set to the minimum water-fat in-phase TE, 2.56 ms for 10.5 T and 2.88 ms for 7 T, for improved sensitivity map estimation. TR was set to 315 ms for 10.5 T and 500 ms for 7 T, and nominal flip angle to 30° for 10.5 T and 38° for 7 T, leading to a total scan time of less than 25 s.

For subsequent analysis of the intrinsic SNR (see the section of “Intrinsic SNR analysis” for its definition), actual flip angle imaging (AFI) (Yarnykh, 2007), measuring the flip angle distribution across the entire brain, was acquired with isotropic 4 mm resolution, 50° nominal flip angle and TR1/TR2 of 20/100 ms.

To facilitate subsequent anatomy-based imaging processing and analysis, whole-brain anatomical reference images were collected using T_1_-weighted (T_1_w) 3D magnetization prepared two rapid acquisition gradient echoes (MP2RAGE) sequence (Marques et al., 2010) at 7 T. For each volunteer, the MP2RAGE was acquired with isotropic 0.7 mm resolution, 2.36 ms TE, 4 s TR, and TI_1_/TI_2_= 740/2430 ms.

To examine the extent of parallel imaging potential at 10.5 T, data were acquired using the highest density (128Rx) RF coil due to its superior SNR and parallel imaging performance (Lagore et al., 2025). For this purpose, the 0.5 mm T_2_* ME-GRE protocol used for the 80Rx coil was applied with higher 2D acceleration factors of 3×3 and 3×4 with CAIPI, in addition to the 2×3 protocol. In all cases, the embedded volumetric navigator images were collected in the same way as described above.

### 2.2 Image reconstruction

All 3D T_2_*w ME-GRE images were reconstructed with in-house MATLAB (Mathworks, Natick, MA, USA) software (https://github.com/jiaen-liu/moco). The reconstruction approach was based on a unified signal model incorporating intra-scan rigid-body motion and B_0_ changes for motion and B_0_ correction, and coil sensitivity maps for SENSE-based parallel imaging reconstruction (Pruessmann et al., 1999). Coil sensitivity maps were estimated from the 2D GRE reference scans using an in-house algorithm, including normalization of individual channel images by the channel-combined image followed by spatial smoothing (Liu et al., 2020). Motion time series was estimated from the navigator magnitude images using an iterative multi-resolution image registration approach (Thevenaz et al., 1998), whereas B_0_ changes over time were estimated based on the navigator phase images. To highlight the importance of joint motion and B_0_ correction, the T_2_*w GRE images were reconstructed using the same data but in three correction modes: (1) motion and spatially linear B_0_ change correction (MoCo+B_0_Co), (2) motion only correction (MoCo) and (3) no correction (NoCo). Details about the implementation of the correction and reconstruction algorithm can be found in our previous publications (Liu et al., 2020; van Gelderen et al., 2023).

### 2.3 *R_2_\** and *χ* quantification

To demonstrate the utility of the 10.5 T ME-GRE data for quantitative mapping of *R_2_\** and *χ*, we analyzed the data acquired with the 80Rx coil. For *R_2_\** mapping, voxel-wise decay rate *R_2_\** values were calculated based on nonlinear least square fitting of the mono-exponential model to the ME-GRE magnitude images. For quantitative susceptibility mapping (QSM), voxel-wise susceptibility *χ* values were quantified using the pipeline implemented in the JHU/KKI QSM toolbox (https://github.com/xuli99/JHUKKI_QSM_Toolbox) (Bao et al., 2016; Li et al., 2019; van Bergen et al., 2016). Briefly, the QSM pipeline included the following processes: path-based phase unwrapping (Abdul-Rahman et al., 2005), brain masking using FSL BET (Smith, 2002), background field removal combining LBV (Zhou et al., 2014) and VSHARP (Wu et al., 2012), and dipole inversion using a modified structural feature collaborative reconstruction approach (Bao et al., 2016) based on a nonlinear data fidelity cost function (Milovic et al., 2018).

### 2.4 Image processing

At the subject level, multi-echo T_2_*w GRE raw images, as well as extracted *R_2_\**, *χ*, and flip angle maps, and MP2RAGE image, were registered to the subject’s 7 T T_2_*w GRE space. This was done by aligning each of these images with the 7 T echo-averaged T_2_*w GRE magnitude image using multi-contrast registration algorithms implemented in the Advanced Normalization Tools (ANTs) (Avants et al., 2008). The registered MP2RAGE was also processed in Freesurfer (Fischl, 2012) to select a layer in cortical grey matter (GM) and another in nearby white matter (WM) from which SNR and *R_2_*\* were sampled to evaluate the intrinsic SNR and precision metrics (see below for details). Specifically, the GM layer was selected to be the mid-layer of the cortical ribbon, whereas the nearby WM layer to be the one in WM tissue that was below the GM-WM border by a distance equal to 40% of the nearby cortical thickness. An illustration of the selected GM and WM layers can be found in the supplemental Fig. S1.

### 2.5 Intrinsic SNR analysis

To underscore the SNR gains with the 10.5 T acquisition relative to its 7 T counterpart, we evaluated intrinsic SNR (iSNR) around the periphery of the cerebrum (Fig. S1) based on the four-echo data acquired with the 80Rx coil and compared the result against that of 7 T. Here, iSNR in unit of 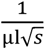 is defined as the SNR of the equilibrium magnetization obtainable in unit volume of medium and unit acquisition time (Edelstein et al., 1986). This metric allows to characterize SNR of the MRI system independent of imaging parameters. To quantify iSNR, the following steps were performed. First, SNR of individual echo images was calculated. The noise map was determined based on the measured noise covariance matrix, sensitivity maps, and the aliasing matrix specific to the parallel imaging scheme employed, as described by Pruessmann et al. (Pruessmann et al., 1999). Here, noise data was acquired with the same receiver bandwidth as the main scan for consistent noise covariance data. Secondly, the SNR at TE=0 (SNR_TE0_) was estimated by applying mono-exponential fitting to the SNR values of the individual echo images. Note that, this can affect the accuracy of iSNR estimation because of the various T_2_* relaxation rates from different water compartments in biological tissue, especially in the WM (Sati et al., 2013). More discussion can be found in the Discussion section. Thirdly, iSNR was computed by dividing SNR_TE0_ by the steady state spoiled GRE signal, to provide SNR measurement independent of the effects of flip angles (obtained from AFI), tissue’s T_1_ relaxation times (obtained from literature) and TR. Finally, the iSNR per unit volume and acquisition time was obtained as divided by the nominal voxel volume and the square root of the total acquisition time for each echo, consistent with methods used in previous studies (Pohmann et al., 2016; Uğurbil et al., 2019). The tissue’s T_1_ relaxation times used were 1.8 s for GM and 1.2 s for WM at 7 T, and 2.1 s for GM and 1.4 s for WM at 10.5 T (Rooney et al., 2007).

### 2.6 *R_2_\** and *χ* precision analysis

To evaluate the precision gains of susceptibility contrast-based quantification at 10.5 T relative to 7 T, we quantified *R_2_\** and *χ* using data acquired with the 80Rx coil and compared the results to those of 7 T. To enable the comparison across different field strengths without the confounding effects of specific sequence parameters and achievable data acquisition duration within each TR, the precision analysis was performed in an idealized condition which assumed instantaneous RF excitation and data acquisition occurring during the entire TR.

To quantify measurement uncertainty, the Cramér–Rao lower bound (CRB) of the standard deviation for *R_2_\** and *χ* estimation was determined. In general, the CRB for an unbiased estimation of a parameter set **θ** from noise-contaminated measurement is derived from the Fisher information matrix **F**, where 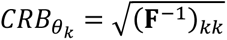 for the *k*th item in **θ** (Assländer et al., 2019; Jones et al., 1996). Here, the elements in **F** are defined as 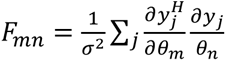 where **y** is the signal as a function of **θ**, 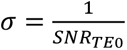 is the standard deviation of noise assuming signal level to be 1 at TE=0, *H* denotes complex conjugate, and *m* and *n* denote indices of matrix elements. Using this framework, the CRB of *R_2_\** 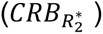 was quantified based on the estimated SNR_TE0_ and *R_2_\** values. *CRB_χ_* was estimated using the relationship 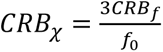, where *f* is the local frequency, *f*_0_ the nominal on-resonance frequency, and the coefficient of 3 reflects the effect of background susceptibility on the frequency distribution inside a sphere (Salomir et al., 2003).

This relationship was confirmed by experimental data in the previous work (van Gelderen et al., 2023) (Fig. 5 in the reference). The CRB of local frequency (*CRB_f_*) was quantified using a linear regression model of the MR signal phase against TE, with the phase signal from each echo weighted by its magnitude to optimize SNR. All CRB data were normalized to unit voxel volume and unit total acquisition time of all echoes.

Due to the field strength dependency of *R_2_\**, the precision of *R_2_\** was evaluated as its CRB normalized by the measured *R_2_*\* difference between GM and WM. By contrast, the precision of *χ* was reported directly as its CRB since it is a field-independent physical quantity if not accounting for field-dependent apparent *χ* due to the underlying microstructure and anisotropic susceptibility effect.

### 2.7 Parallel imaging performance

Parallel imaging performance of all three RF coils under consideration was compared by calculating g-factors for different 2D acceleration factors. With each coil, g-factors were evaluated for five acceleration factors of 2×2, 2×3, 3×3, 3×4, and 4×4, all with 2D CAIPI. In each case, g-factors across the entire brain were quantified as the ratio of the SENSE-based noise map without acceleration over that with acceleration and further normalized by 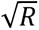 where *R* is the total combined acceleration factor (Pruessmann et al., 1999), using the estimated sensitivity maps from the multi-slice 2D GRE reference scan. Furthermore, inverse g-factor (1/*g*) values measuring the retained SNR were calculated and their statistics, including the median values across the entire brain, were derived to evaluate parallel imaging performances.

## 3. Results

The reconstruction with simultaneous motion and B_0_ correction (i.e., MoCo+B_0_Co) produced the best image quality (Fig. 1), effectively eliminating image artifacts observed under the other two reconstruction modes. Although outperforming the reconstruction without any correction, the reconstruction with motion correction alone (MoCo) still showed image artifacts especially for the longer TE (∼25 ms).

**Figure 1.**
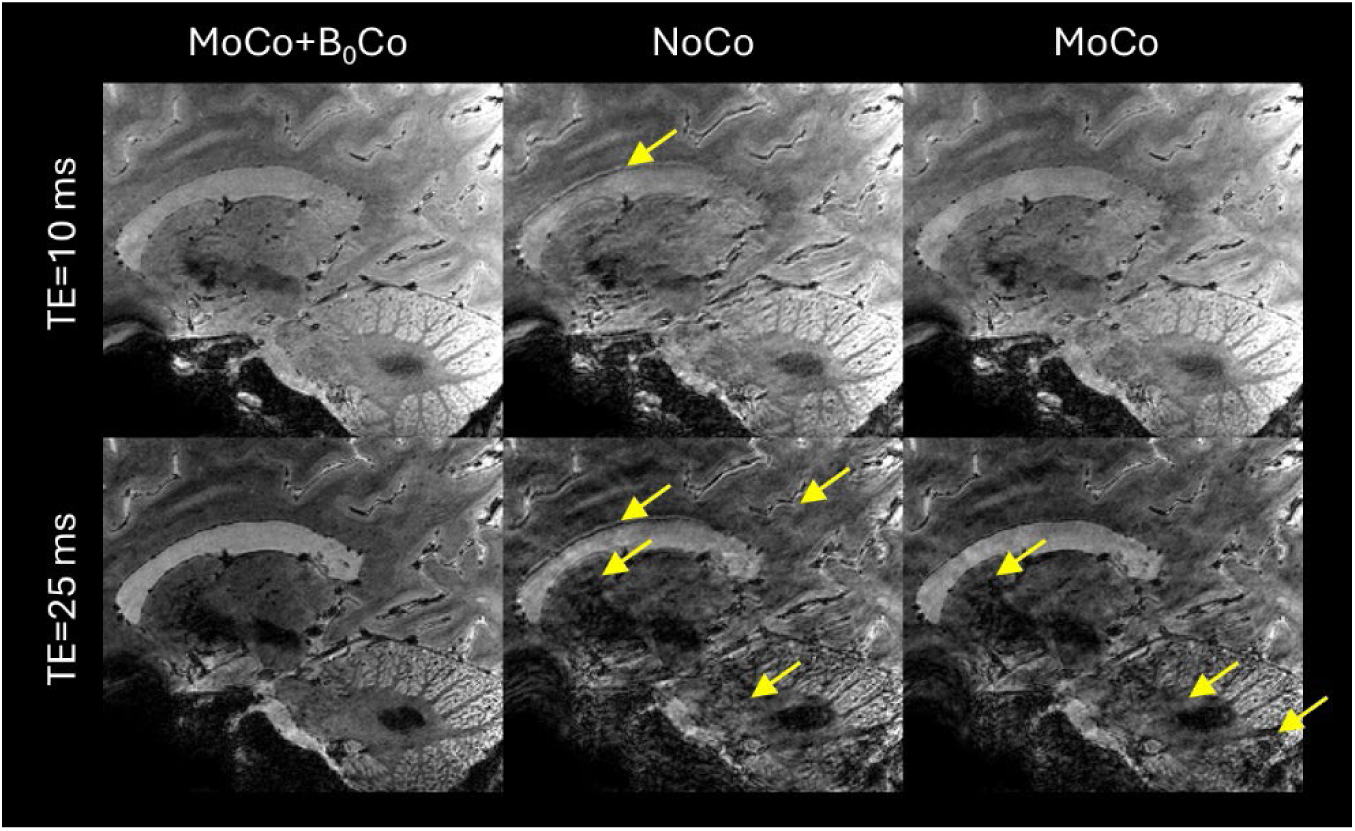
Importance of simultaneous motion and B_0_ corrections in preserving the image quality for *T_2_\**-weighted multi-echo 3D gradient echo (GRE) brain MRI at 10.5 T. Shown are *T_2_\**-weighted GRE magnitude images acquired with isotropic 0.5 mm resolution and the 80Rx coil in a sagittal slice at two echo times of ∼10 ms (top) and ∼25 ms (bottom). Images were reconstructed with simultaneous motion and B_0_ correction (MoCo+ B_0_Co), with no correction (NoCo), and with only motion correction (MoCo), all based on the same data. Note how the reconstruction with simultaneous motion and B_0_ corrections produced the best image quality, effectively eliminating image artifacts especially for later echoes. Arrows point to artifacts.

The ME-GRE image reconstruction with simultaneous motion and B_0_ correction translated into high quality quantitative *R_2_*\* and *χ* maps across the entire brain (Fig. 2), which in turn, enabled clear visualization of fine-scale brain structures, e.g., the hippocampal superficial medullary stratum (Adachi et al., 2003) (Fig. 3), a fiber-rich layer receiving inputs from the entorhinal cortex and deep nuclei.

**Figure 2.**
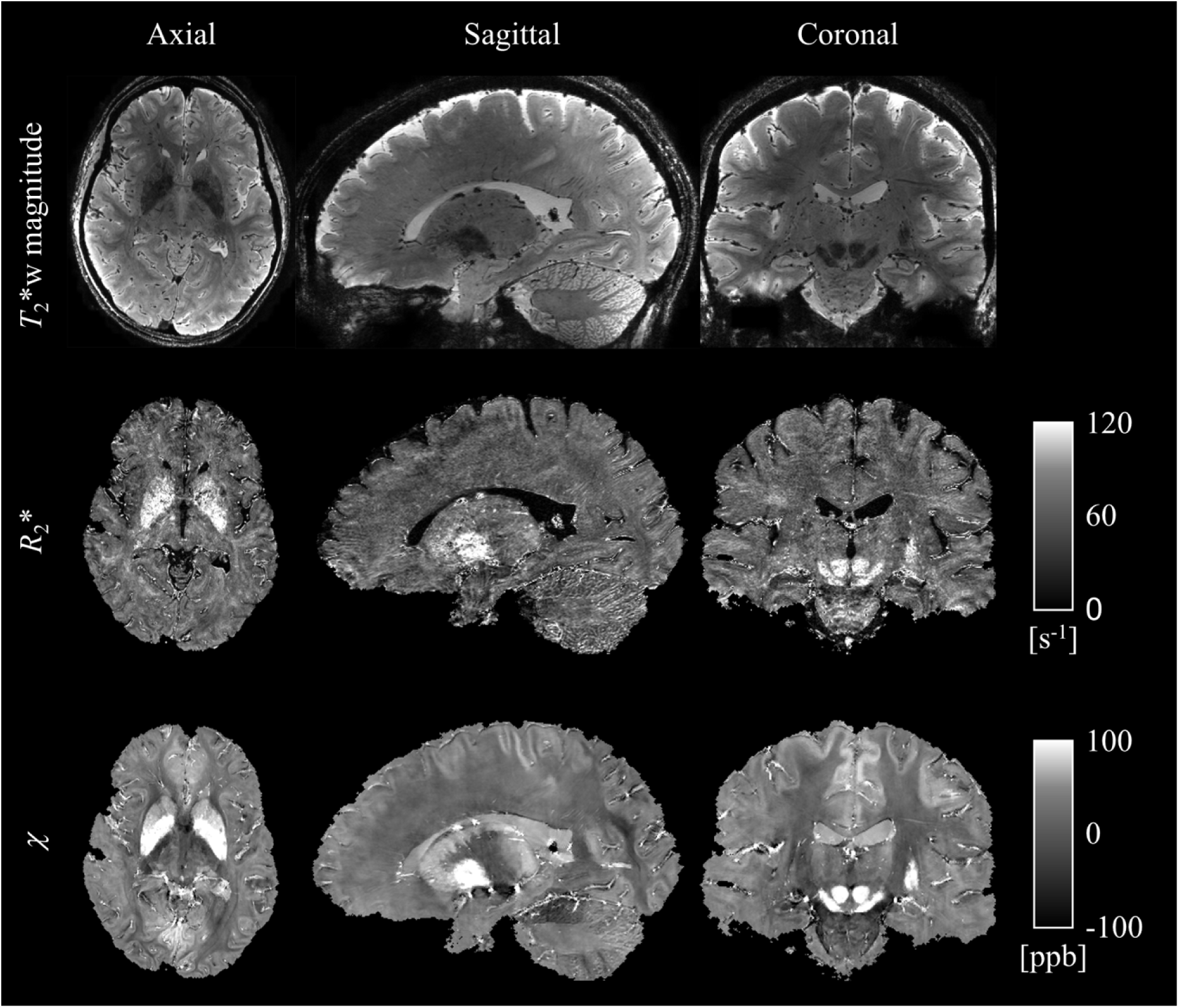
Utility of the motion-robust *T_2_*\*-weighted multi-echo GRE MRI for whole brain multi-parametric mapping at 10.5 T. Shown are echo-averaged magnitude images (top), decay rate *R_2_*\* maps (middle) and magnetic susceptibility *χ* maps (bottom) in three orthogonal views in the same healthy volunteer as in Fig. 1, with 0.5 mm resolution and reconstructed with simultaneous motion and B_0_ correction. The results demonstrated high quality multi-echo GRE images and the corresponding parameter mapping across the entire brain.

**Figure 3.**
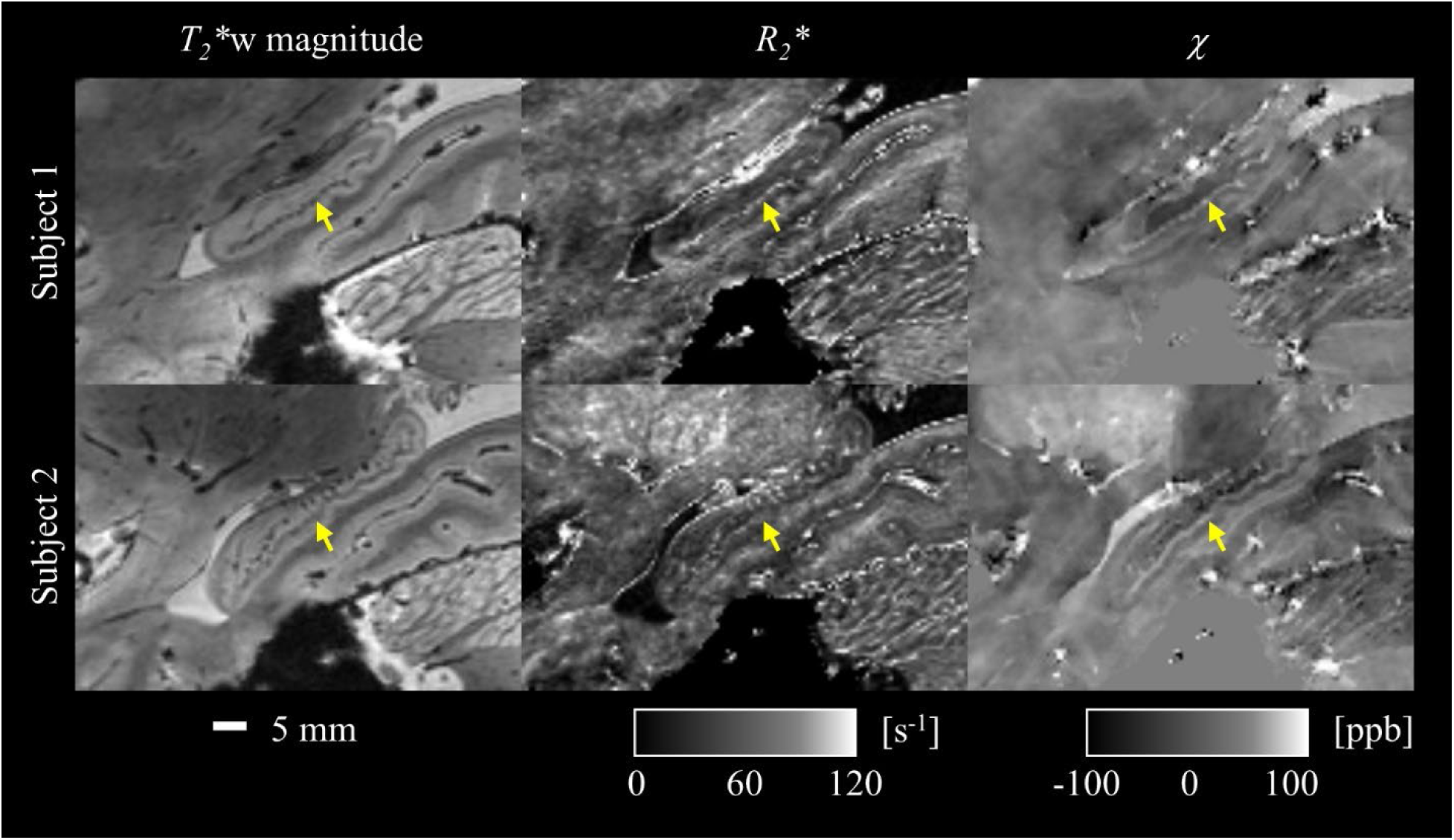
Demonstration of the capability of the utilized motion-robust *T_2_\**-weighted multi-echo GRE MRI at 10.5 T in delineating fine scale brain structures. Shown are zoomed-in sagittal views of the *T_2_\**-weighted echo-averaged magnitude, *R_2_*\* and *χ* images focused on the hippocampus in two healthy subjects. The high-resolution images demonstrated clear visualization of the internal structures in hippocampus, including its fiber-rich superficial medullary stratum (as marked by arrows).

Compared to 7 T, 10.5 T allowed substantial SNR and precision gains (Table 1 and Fig. 4). The subject averaged iSNR gain across the periphery of the cerebrum (Table 1) improved by 42% for GM and 36% for WM at 10.5 T. As shown in Fig. 4, the improvement is more noticeable in the parietal and occipital regions. Using these iSNR values and the calculated tissue-specific *R_2_*\* results (Table 2), the precision analysis indicates improved precision for both *R_2_\** and *χ* quantification (Figs. 5 C-F). Again, due to the field dependence of *R*_2_*, the 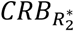 was normalized by the subject-averaged GM-WM *R_2_\** difference (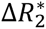, 7.5 s^−1^ at 10.5 T and 5.5 s^−1^ at 7 T) to highlight the precision of *R_2_\** in differentiating GM and WM (Figs. 5 C-D). In both subjects, the ratio between the WM-GM *R_2_*\* difference and mean *R_2_\** is similar at 10.5 T and 7 T (Table 2).

**Figure 4.**
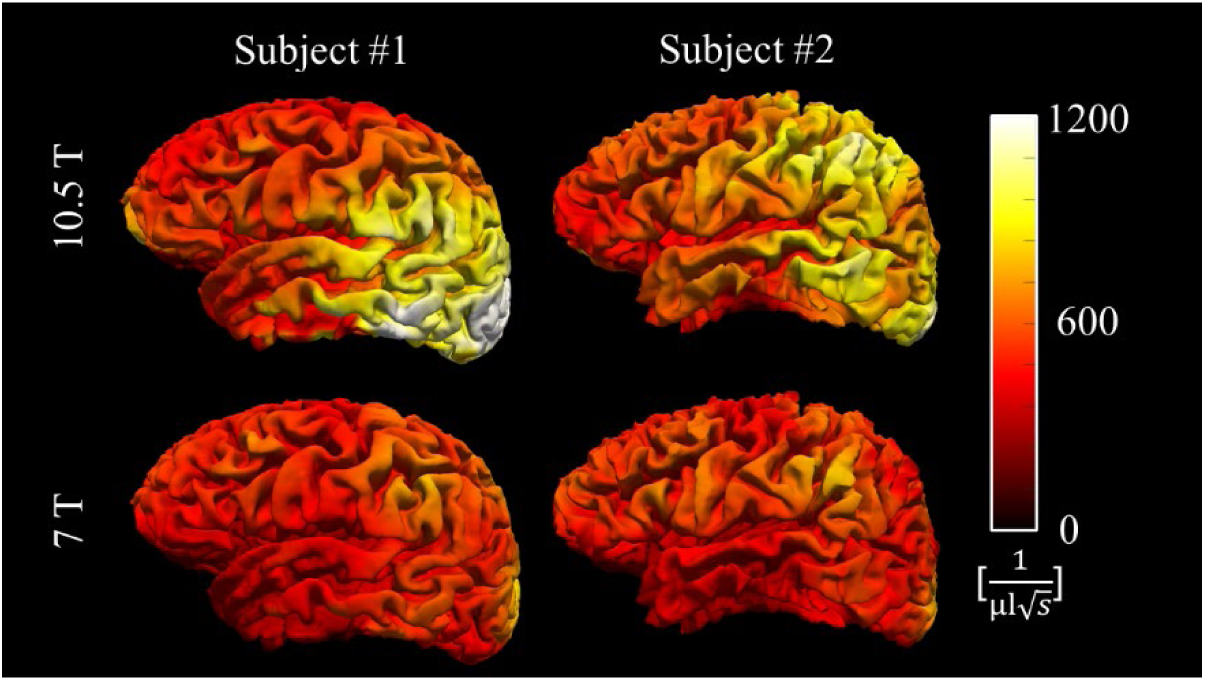
Intrinsic SNR improvement obtained in the cortical gray matter. ISNR distributions are shown in the left hemisphere for both subjects at 10.5 T and 7 T.

**Figure 5.**
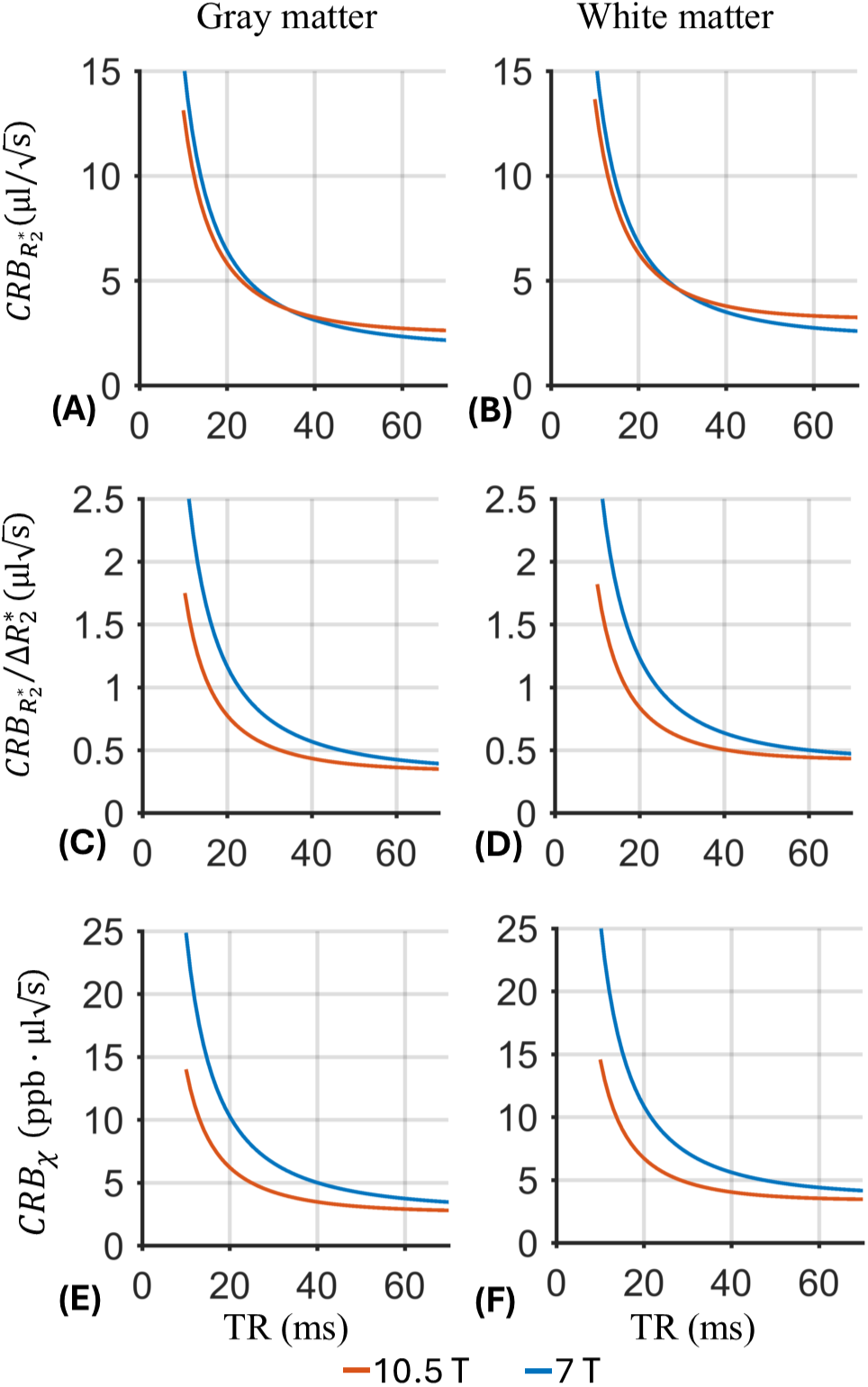
Improved precision efficiency for *R_2_\** and *χ* quantification using the 80Rx coil at 10.5 T vs. 32Rx coil at 7 T in gray and white matter, respectively. Shown are the uncertainty of *R*_2_* as measured by its Cramér–Rao lower bound (CRB) (A and B), the normalized 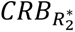 by the difference of *R_2_\** between gray and white matter at specific field strength (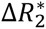, 7.5 s^−1^ at 10.5 T and 5.5 s^−1^ at 77 T) (C and D), and the uncertainty (CRB) of *χ* (E and F). All data were normalized to unit voxel volume and unit total scan time. Note that the results demonstrate dependency on the excitation repetition time (TR) assuming data acquisition occurred during the entire TR.

**Table 1.** Intrinsic SNR (iSNR, unit: 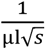) measured in the selected cortical layer (GM) and adjacent white matter layer (WM) across the entire peripheral cerebrum at 10.5 T and 7 T. The iSNR gain was calculated as the mean of location-wise gain ratio in the GM and WM layers, respectively. An example of the GM and WM layers can be found in Fig. S1.

|  | 10.5 T |  | 7 T |  | iSNR gain |  |
| --- | --- | --- | --- | --- | --- | --- |
|  | GM | WM | GM | WM | GM | WM |
| Subject #1 | 677 | 549 | 495 | 418 | 1.41 | 1.34 |
| Subject #2 | 678 | 565 | 494 | 423 | 1.42 | 1.38 |

**Table 2.** *R_2_\** (s^−1^) measured at 10.5 T and 7 T for gray (GM) and white matter (WM) and their relative contrast 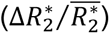. Here, 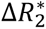 denotes the location-wise *R_2_\** difference between individual pairs of adjacent WM and GM locations from the corresponding layers, and 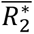 represents their mean. 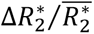 was averaged along the selected GM and WM layers as illustrated in Fig. S1.

|  | 10.5 T |  |  | 7 T |  |  |
| --- | --- | --- | --- | --- | --- | --- |
| | GM | WM | $\Delta R_2^*/\overline{R_2^*}$ | GM | WM | $\Delta R_2^*/\overline{R_2^*}$ |
| Subject #1 | 47 | 53 | 0.13 | 33 | 37 | 0.12 |
| Subject #2 | 42 | 51 | 0.20 | 29 | 36 | 0.21 |

Both high-density RF coils at 10.5 T outperformed the 7 T Nova 32Rx coil in parallel imaging performance (Fig. 6), retaining SNR gain as measured by the inverse g-factor (1/*g*) especially with higher acceleration factors. For relatively low acceleration factors of 2×2 and 2×3, all three RF coils presented similar parallel imaging performances with comparable 1/*g* values across the brain. However, for higher acceleration factors, the 10.5 T high-density coils demonstrated significantly better parallel imaging performance. For example, in comparison with the 7 T 32Rx coil, the median value of 1/*g* across the brain using the 128Rx coil increased by 3.2% (0.96 vs. 0.93) for the acceleration factor 3×3, by 6.9% (0.93 vs. 0.87) for 3×4, and by 22.5% (0.87 vs. 0.71) for 4×4. Moreover, at 10.5 T, the 128Rx coil outperformed its 80Rx counterpart in parallel imaging performances, increasing the 1/*g* median value by 3.2% (0.96 vs. 0.93) for 3×3, by 3.3% (0.93 vs. 0.90) for 3×4, and by 2.4% (0.87 vs. 0.85) for 4×4. Accordingly, the use of the 10.5 T 128Rx coil enabled data collection with a total acceleration factor of 12 (Fig. 7), giving rise to visually comparable image quality to those obtained with lower acceleration factors.

**Figure 6.**
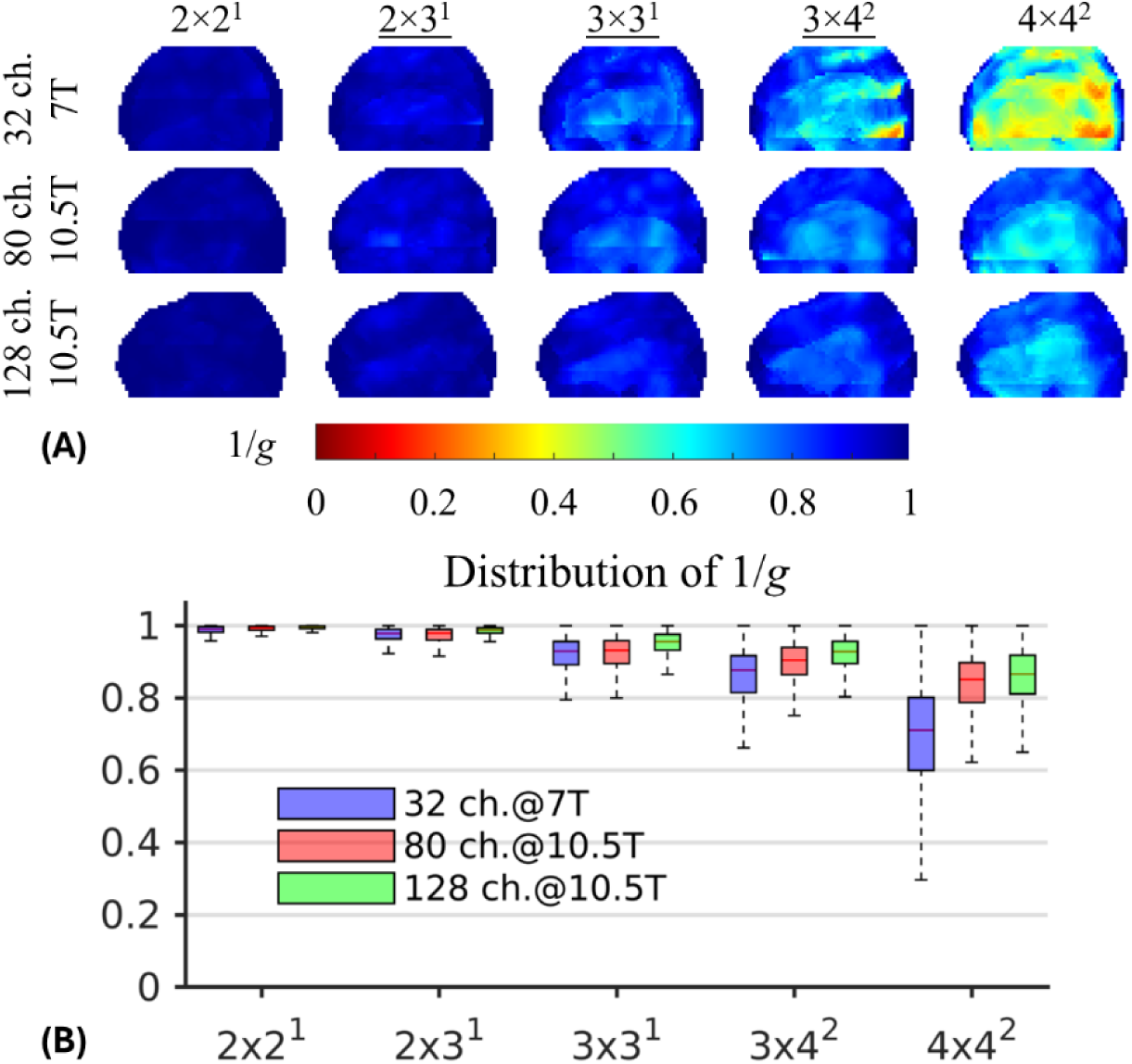
Improved parallel imaging performance using the high-density 10.5 T coils (one with 80 receive channels and another with 128 receive channels) vs. the commercial 7 T Nova 32-channel coil. For each coil the minimum intensity projections of the inverse g-factor (1/*g*) maps in the sagittal view are shown for different 2D acceleration schemes (A), alongside the corresponding boxplots characterizing the statistical distribution of 1/*g* across the entire brain (B). Here, the 2D acceleration schemes are denoted as *R*_p_ × *R*_s_ ^Δs^ where *R*_p_ and *R*_s_ stand for the acceleration rate in the phase (left-right) and slice (head-foot) directions, respectively; Δs is the phase encoding shift in the slice direction for controlled aliasing. For boxplots, the horizontal bar within each box indicates the median value and the box height represents the 25th to 75th percentile range. Note that the relative parallel imaging improvement of the 10.5 T coils compared to the 7 T 32-channel coil was more prominent with higher acceleration factors.

**Figure 7.**
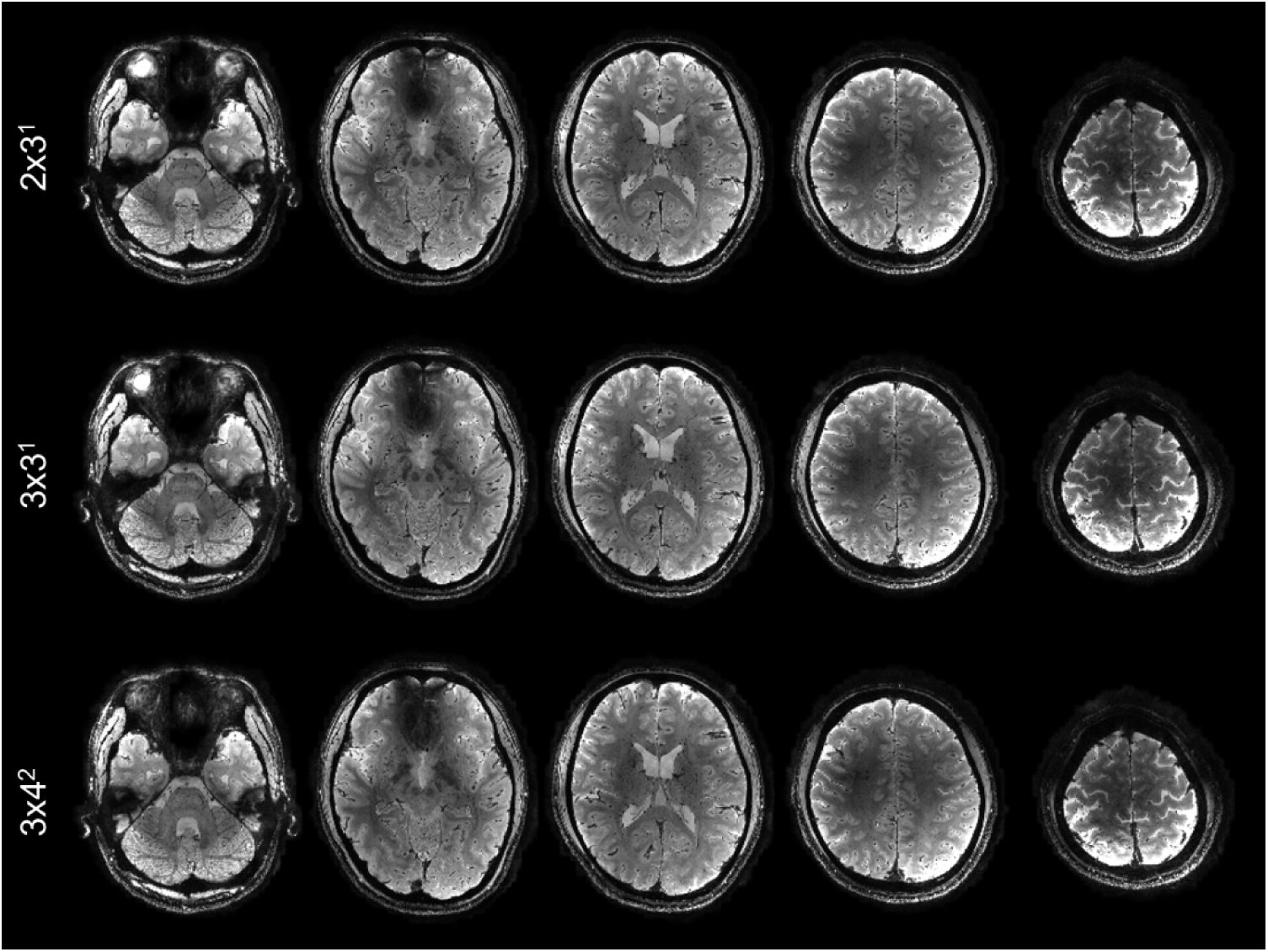
Highly accelerated whole-brain T_2_*-weighted multi-echo GRE MRI at 10.5 T, acquired using the 128-channel RF receive array. Shown are T_2_*-weighted echo-averaged magnitude images in five representative axial slices with different 2D acceleration schemes. All images were reconstructed with simultaneous motion and B_0_ corrections at isotropic 0.5 mm resolution from the same subject. Even with a combined 12-fold acceleration, the overall imaging approach was able to retain the image quality.

## 4. Discussion

We have demonstrated the feasibility of mesoscopic whole-brain multi-echo T_2_*-weighted anatomic MRI at 10.5 T and its utility for multi-parametric brain mapping in living humans. Critical to the success was a synergistical combination of technical developments including high-density RF coil design and motion-robust 3D ME-GRE image acquisition. With custom 80-channel and 128-channel RF coils and navigator-based motion-B_0_-corrected image reconstruction, whole brain imaging at 0.5 mm isotropic resolution was demonstrated. The results (Figs. 1-3) showed that image reconstruction with joint motion and B_0_ correction resulted in high-quality multi-echo T_2_*w images, which in turn led to high-quality *R_2_*\* and *χ* mapping, delineating fine brain structures. Further comparison against data collected at 7 T using the commercial Nova 32-channel RF coil suggested that the 10.5 T setup can yield substantial SNR and precision gains (Table 1 and Fig. 5). Evaluation of parallel imaging performances (Fig. 6) indicated that high-density coils at 10.5 T, especially the 128-channel coil, substantially reduced g-factors compared to the 7 T setup with the Nova coil, allowing for high-fidelity image reconstruction with total acceleration factors greater than 10 (Fig. 7).

### 4.1 Motion and B_0_ correction

As shown in the current work (Fig. 1) and our previous work (van Gelderen et al., 2023), even subtle motion can introduce substantial artifacts surrounding mesoscopic structures. Motion (including head and torso motion) also introduces B_0_ field fluctuation which affects T_2_*w image quality (Liu et al., 2018). Unlike our previous work where correcting the spatially nonlinear B_0_ changes was found to be effective in reducing image artifact when the subject was instructed to move the head (Liu et al., 2020), here, only linear B_0_ correction was performed given the moderate head motion (Fig. S2). In our previous work including more subjects (n=11) and longer scan times (∼ 35 min) (van Gelderen et al., 2023), the result suggested that nonlinear B_0_ correction provided limited improvement in the moderate head motion regime but at the cost of long reconstruction time. For more severe head motion, other factors in addition to more severe nonlinear B_0_ changes, such as transmit and receive B_1_ inconsistency, large gaps in k-space and limited temporal resolution and motion and field measurement accuracy of the current navigator design, can contribute to image artifacts. The joint effect of these different sources may warrant future investigation.

### 4.2 Gain of SNR and precisions of *R*_2_* and *χ* quantification at 10.5 T

The intrinsic SNR analysis (Table 1) shows that the peripheral iSNR improved by as much as 42% from 7 T (Nova 32Rx) to 10.5 T (80Rx). Although this result was obtained utilizing the accelerated 0.5 mm GRE data with a 2×3 acceleration factor, given the minimal and similar g-factor penalty for both coils within this acceleration regime (Fig. 6), the iSNR gain is expected to be similar to the result without acceleration and indeed agreed with results obtained in previous human and phantom experiments (Waks et al., 2025).

The 10.5 T system is a new and “immature” system that has not experienced the refinements that exists in contemporary 7 T instruments and 7 T RF coils. It has proven, for example, difficult to capture the ultimate intrinsic SNR (uiSNR) in the human head at 10.5 T using RF coil designs inspired by approaches used at 7 T and lower magnetic fields (Lagore et al., 2025; Waks et al., 2025). The coils employed in this study were shown to perform as efficiently as simpler 7 T coils (e.g the 32Rx coil employed in this study) in capturing uiSNR in the center, albeit with much greater complexity in channel number and layout; however, the periphery of the brain where uiSNR is in principle extremely high may be captured with very low efficiency even with the 128 channel coil employed in this study. As such the results at 10.5T should be regarded as preliminary documentation of gains available at this very high magnetic field which will only improve as new coils are developed.

Combined with the observed peripheral iSNR increase (Table 1) and faster T_2_* relaxation at 10.5 T, the results for *R_2_*\* and *χ* (Fig. 5) suggest higher precision efficiency, i.e., precision per unit scan time, at 10.5 T for the same TR applied at both field strengths. With longer TR, the more complete T_2_* relaxation process can be sampled at 7 T relative to 10. 5 T, allowing closer precision assessment between the two field strengths. On the other hand, without losing the precision efficiency, scan time can be reduced at 10.5 T by prescribing field-specific TRs which scale with the T_2_* value. As shown in Figs. 5 C-F, for similar precision metrics, scan time can be reduced by about 1/3 by reducing TR, e.g., utilizing TR=30 ms at 10.5 T vs. TR=45 ms at 7 T, or with TR=20 ms at 10.5 T vs. TR=30 ms at 7 T.

While the above analysis applies to quantification of *R_2_\** and *χ* from ME-GRE signal, the result also sheds light on the CNR gain of T _2_*w signal from 7 T to 10.5 T, such as in T_2_*w anatomical or blood-oxygenation-level-dependent (BOLD) functional imaging applications. With TR<<T_1_ and TR (limited by optimal TE) scaled to field specific T_2_* values and assuming Ernst angles, the CNR efficiency of T_2_*w signal (i.e., CNR per unit scan time) is proportional to 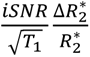 (Duyn, 2012). From Table 2, it can be observed that 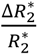 is approximately the same between 7 T and 10.5 T. This leaves the optimal T_2_*w CNR gain to be dominated by the iSNR gain at 10.5 T (e.g., 42% gain in the GM when using the 80Rx coil as observed in this study) and slightly reduced by the increased 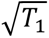 (T_1_=2.1 s at 10.5 T vs. 1.8 s at 7 T). Note that for BOLD signal, a quadratic field-dependency of 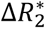 may be observed within blood water or tissue water in the capillary bed sufficiently close to the source of magnetic susceptibility changes, e.g., hemoglobin (Uludağ et al., 2009), rendering the overall CNR gain at 10.5 T to be higher than the prediction based on the reported GM-WM 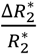 values in Table 2.

### 4.3 Utility of 10.5 T MRI for mesoscopic imaging

With the high SNR and strong susceptibility effect at 10.5 T, the T_2_*w MRI and its quantitative products of *R_2_*\* and *χ* can be exploited for obtaining a better understanding of cortical organization and function in healthy humans. For example, the hyperintense *R^2^*\* and *χ* values as observed in the fiber-rich hippocampal superficial medullary stratum (Adachi et al., 2003) (Fig. 3) indicate high iron concentration, similar to other fiber-rich structures such as the bands of Baillarger in the cortex and the superficial white matter near the GM-WM junction (Fukunaga et al., 2010; Kirilina et al., 2020; van Gelderen et al., 2023). High iron concentration has been reported in oligodendrocytes (Reinert et al., 2019), the main type of cells responsible for generating and maintaining myelin in the brain. Such tissue properties can provide histologically defined information at the subject level to refine in vivo segmentation of mesoscopic structures, such as cortical laminae which are currently determined using geometric segmentation criteria for in vivo applications (Huber et al., 2021; Waehnert et al., 2014).

### 4.4 Limitation of iSNR and *R_2_*\* estimation based on mono-exponential fitting in the white matter

The usage of mono-exponential model in fitting the *R_2_*\* and iSNR in tissue, especially in the WM, can lead to bias due to the underlying heterogeneous relaxation behavior of water molecules in different tissue compartments (Sati et al., 2013). Using relatively long TEs in this study, mono-exponential fitting can underestimate fast decaying components with *R_2_*\* on the order of 100-200 s^−1^ (Sati et al., 2013). This is evident in our results, showing lower iSNR at both field strengths and slightly lower iSNR gain in the WM compared to the GM (Table 1). The latter can be caused by the higher fraction of fast decaying compartments at 10.5 T. Another factor contributing to the apparently low iSNR in the WM is the abundant lipid macromolecular proton in the WM, whose extremely short T_2_ on the order of 10-100 μs renders them undetectable at the current TEs (Wilhelm et al., 2012; van Gelderen et al., 2017).

### 4.5 Limitation related to RF pulse design

One limitation of the current study was that all data collection at 10.5 T was performed with RF excitation in the CP mode, mimicking a conventional birdcage-type single-channel transmit setup. This led to RF shading artifacts observed across the brain especially in upper and lower brain regions arising from RF inhomogeneity. Also related to RF pulses, using a traditional binomial RF pulse for water-selective excitation resulted in a ring-shaped signal dropout around the lower frontal lobe where relatively large off-resonance existed. This pulse was adopted to minimize the fat signal contamination that would otherwise degrade the reconstruction of the EPI-based navigator images. One effective solution to both the RF shading artifact and ring-shaped signal dropout is to use parallel transmission (pTx) (Katscher et al., 2003; Padormo et al., 2016). Part of our future work is to combine the custom high-density RF coils (both allowing for 16-channel pTx) with intelligent pTx spatial spectral pulse design (Shao et al., 2025) to achieve uniform water-selective excitation across the entire brain even in the presence of large off-resonance. This will be important to realize the full potential of 10.5 T, including high SNR, T_2_* contrast and parallel imaging performance as shown here, in mesoscale motion-robust whole-brain T_2_*w imaging.

### 4.6 Imaging speed gain through 3D EPI

In this study, we showcased the ability to acquire quality mesoscopic whole brain T_2_*w anatomical MRI with isotropic 0.5 mm resolution at 10.5 T using a 3D ME-GRE sequence. With a combined 6-fold acceleration, the total scan time was ∼13.3 min even with a very short TR of 35 ms. However, for T_2_*w MRI at even higher resolution, the use of a 3D GRE sequence starts to fall short of acquisition efficiency, potentially leading to prohibitively long scan times. For even higher resolutions in the range of 0.3 – 0.4 mm, it is recommended that a 3D multi-shot multi-echo EPI sequence should be used, given its demonstrated ability to effectively reduce the total scan time without compromising image quality (Huang et al., 2023; Zwanenburg et al., 2011). With the 3D EPI sequence, multiple GRE readout lines, the number of which is known as the EPI factor, can be collected for each echo time per TR (as opposed to a single readout line for each echo time per TR in the 3D GRE sequence). The EPI factor serves as an additional acceleration factor without parallel imaging penalty. Recently, our own work has demonstrated the utility of 3D EPI for isotropic 0.4 mm resolution multi-echo T_2_*w MRI at 10.5 T, with whole brain and cerebellum focused coverage in about 10 minute scan time (Qu et al., 2025b, 2025a). Part of our future work is to investigate the optimal strategy of combining our motion-robust 3D EPI method with denoising (Ye et al., 2025) to ensure image quality at unprecedented isotropic 0.3 mm resolution for multi-echo T_2_*w MRI.

### 4.7 Navigator design

The 3D EPI volumetric navigator utilized in the ME-GRE sequence can be improved in the context of T_2_*w MRI at UHF. First, instead of using the short TE before the high resolution GRE acquisition, acquiring the navigator at longer TE can improve the quantification of *R_2_*\* and *χ* given the much faster T_2_* decay at 10.5 T. In our previous work, we had illustrated the feasibility and accuracy of a longer TE navigator for motion and B_0_ correction despite of lower navigator SNR and phase wrapping issues (van Gelderen et al., 2023). Secondly, alternative k-space trajectories that cover 3D space more efficiently and facilitate higher acceleration factors can be explored for volumetric navigator acquisition to minimize the navigator acquisition time or improve its temporal resolution, such as the spherical stack of spirals (Assländer et al., 2013) and rotating radial EPI approaches (Rettenmeier et al., 2022).

## 5. Conclusion

We demonstrated the feasibility to perform mesoscopic whole-brain T_2_*w anatomical MRI in humans at 10.5 T. Using custom high-density RF coils in combination with a motion-robust multi-echo 3D GRE sequence, we have presented data showing superior SNR, evaluated the susceptibility contrast and assessed the parallel imaging performance at 10.5 T in comparison with 7 T. At 10.5 T, high-quality whole-brain T_2_*w images and quantitative *R_2_*\* and *χ* mapping at isotropic 0.5 mm resolution were obtained, delineating fine-scale brain structures. This work paves the way for future applications aimed at studying neuroanatomy in humans with resolutions beyond what is attainable with existing imaging approaches.

## Acknowledgement

The authors would like to honor and remember Pierre-Francois Van de Moortele for his invaluable contributions to the development of ultrahigh field MRI and the 10.5 T MRI. The authors would like to thank Jerahmie Radder, Scott Lynch, and Carlos Soria from the CMRR for their assistance with setting up computation resources. This work was supported in part by Hamon Foundation, Texas Instrument Foundation, NIH grants (R01 NS136490, P41 EB027061, U01 EB025144, and S10 RR029672), and the intramural research program of the National Institute of Neurological Disorders and Stroke at NIH.

## Author Contributions

**Jiaen Liu:** Conceptualization, Funding acquisition, Methodology, Formal analysis, Writing – original draft, Writing – review & editing

**Peter van Gelderen:** Methodology, Writing – review & editing

**Jacco A. de Zwart:** Methodology, Writing – review & editing

**Jeff H. Duyn:** Funding acquisition, Methodology, Writing – review & editing

**Yujia Huang:** Methodology

**Andrea Grant:** Methodology

**Edward Auerbach:** Methodology

**Matt Waks:** Methodology, Writing – review & editing

**Russell Lagore:** Methodology, Writing – review & editing

**Lance Delabarre:** Methodology

**Alireza Sadeghi-Tarakameh:** Methodology, Writing – review & editing

**Yigitcan Eryaman:** Methodology, Writing – review & editing

**Gregor Adriany:** Methodology, Writing – review & editing

**Kamil Ugurbil:** Funding acquisition, Methodology, Writing – review & editing

**Xiaoping Wu:** Conceptualization, Funding acquisition, Methodology, Writing – original draft, Writing – review & editing

## Data and Code Availability

Data can be made available upon request subjective to funding agency and institute policies.

## Supplementary Figures

**Fig. S1.**
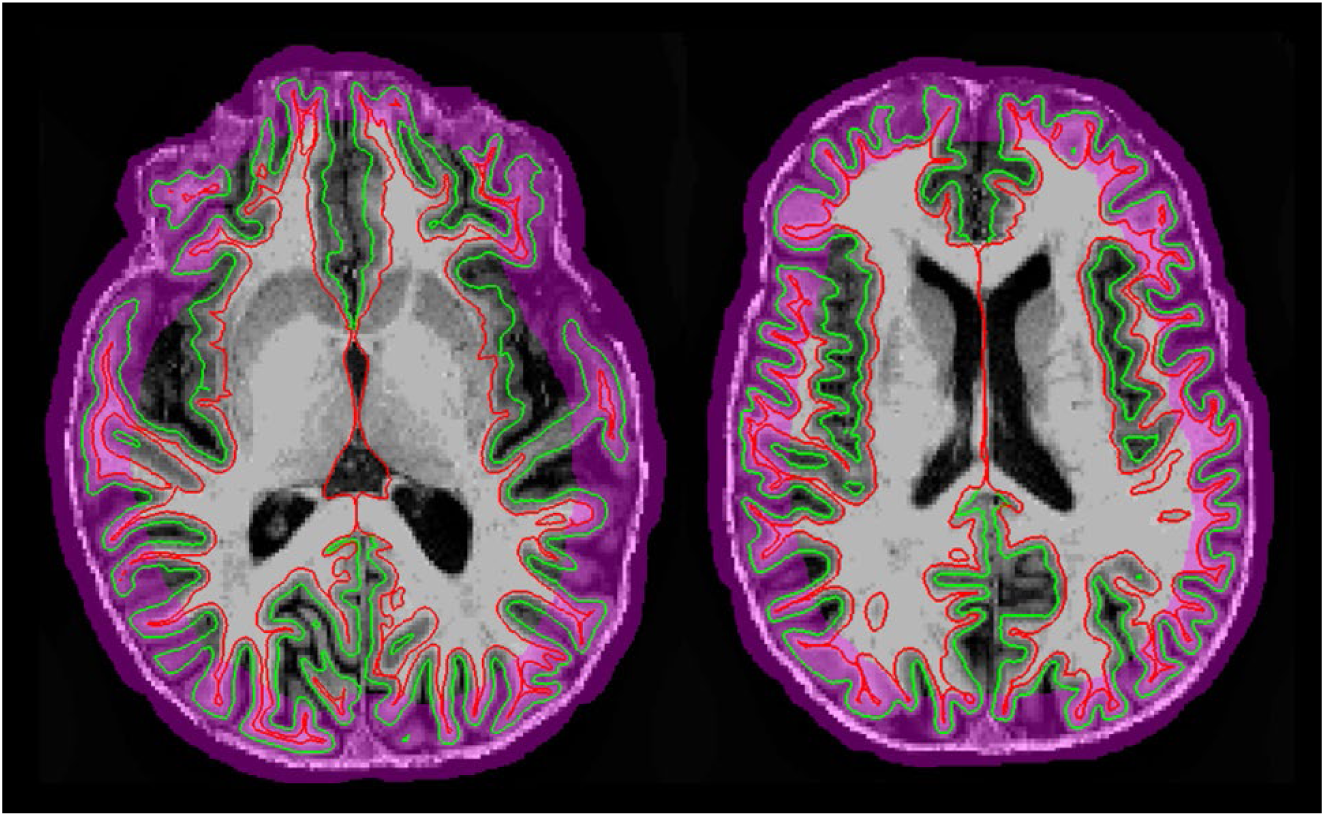
Regions included for intrinsic SNR and *R_2_*\* contrast analysis near the periphery of the cerebrum as defined by the purple band. Two representative axial slices from one volunteer are shown as examples. Data analysis was performed within the purple band, along the selected cortical gray matter layer (as indicated by the green contour), and its adjacent white matter layer (as indicated by the red contour).

**Fig. S2.**
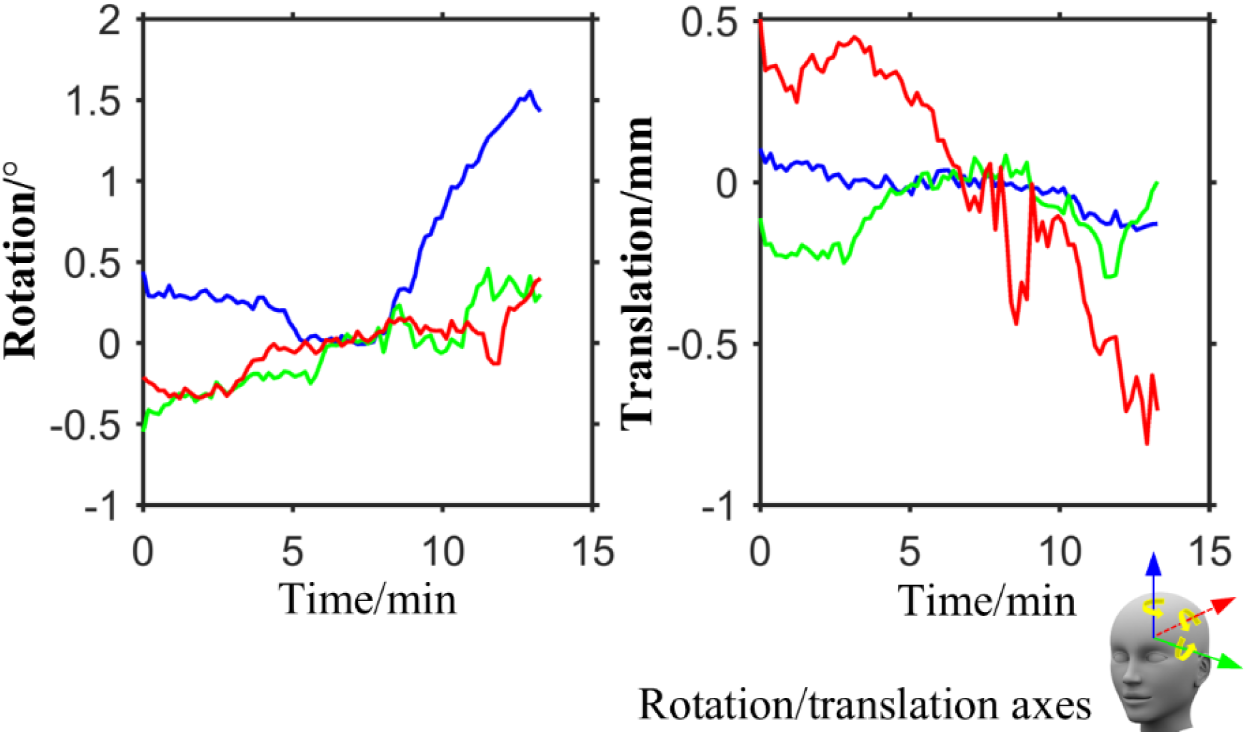
Motion tracking based on navigators. Shown are the navigator-measured rotation and translation time courses in one volunteer during the scan. The three axes that define the rotation and translation are color coded as illustrated in the human head model on the bottom right. Note that moderate head motion was observed throughout the ∼13.3 min 10.5 T scan associated with Fig. 1 in the main text.

