## Supplemental Figures for "Mesoscopic whole-brain T_2_*-weighted and associated quantitative MRI in healthy humans at 10.5 T"

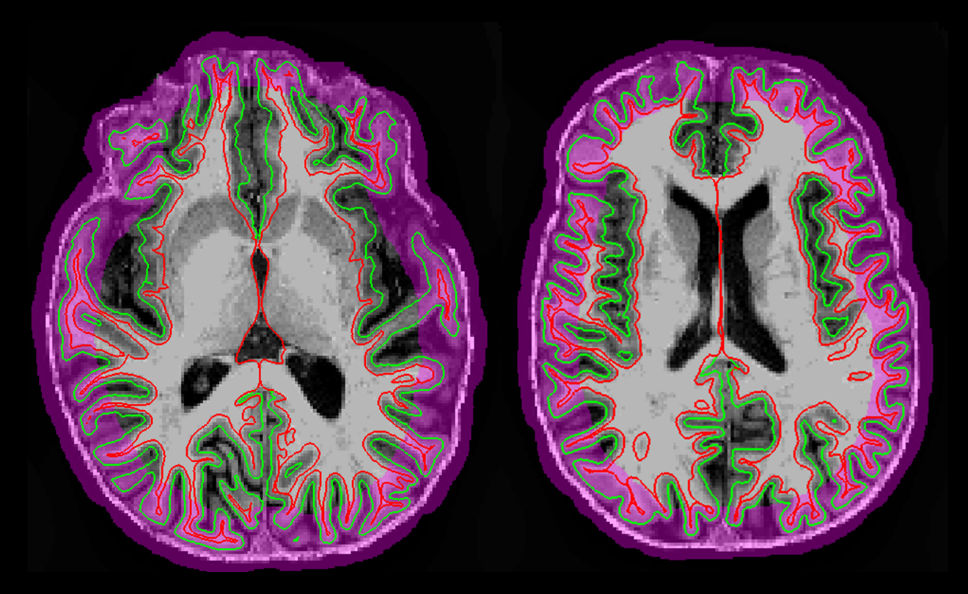

Fig. S1. Regions included for intrinsic SNR and *R*_2_^*^ contrast analysis near the periphery of the cerebrum as defined by the purple band. Two representative axial slices from one volunteer are shown as examples. Data analysis was performed within the purple band, along the selected cortical gray matter layer (as indicated by the green contour), and its adjacent white matter layer (as indicated by the red contour).

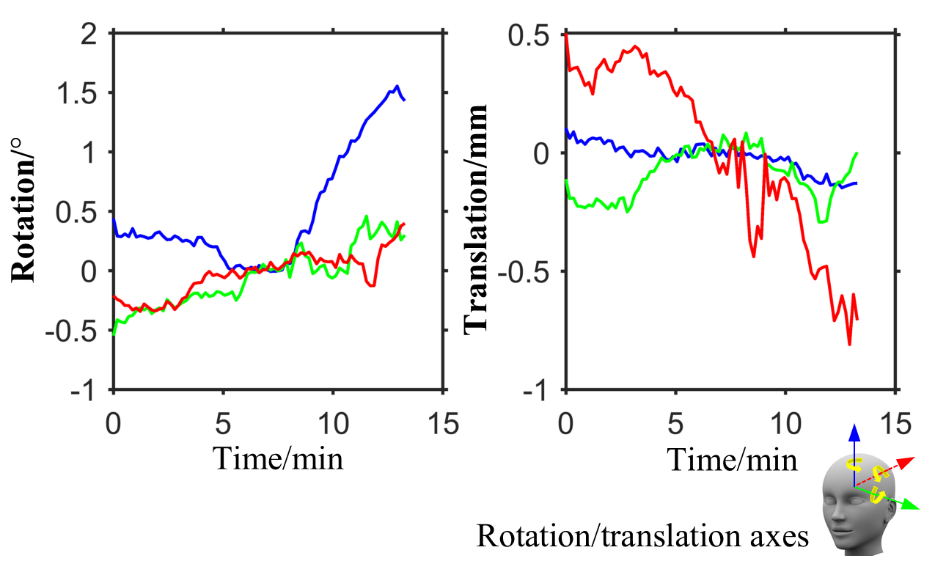

Fig. S2. Motion tracking based on navigators. Shown are the navigator-measured rotation and translation time courses in one volunteer during the scan. The three axes that define the rotation and translation are color coded as illustrated in the human head model on the bottom right. Note that moderate head motion was observed throughout the ~13.3 min 10.5 T scan associated with Fig. 1 in the main text.
